# Learning patterns of HIV-1 co-resistance to broadly neutralizing antibodies with reduced subtype bias using multi-task learning

**DOI:** 10.1101/2023.09.28.559724

**Authors:** Aime Bienfait Igiraneza, Panagiota Zacharopoulou, Robert Hinch, Chris Wymant, Lucie Abeler-Dörner, John Frater, Christophe Fraser

**Affiliations:** Pandemic Sciences Institute, Nuffield Department of Medicine, University of Oxford, Oxford, UK; Big Data Institute, Li Ka Shing Centre for Health Information and Discovery, Nuffield Department of Medicine, University of Oxford, Oxford, UK; Peter Medawar Building for Pathogen Research, Nuffield Department of Medicine, University of Oxford; NIHR Oxford Biomedical Research Centre, Oxford University Hospitals NHS Foundation Trust, John Radcliffe Hospital, Oxford, UK

## Abstract

The ability to predict HIV-1 resistance to broadly neutralizing antibodies (bnAbs) will increase bnAb therapeutic benefits. Machine learning is a powerful approach for such prediction. One challenge is that some HIV-1 subtypes in currently available training datasets are underrepresented, which likely affects models’ generalizability across subtypes. A second challenge is that combinations of bnAbs are required to avoid the inevitable resistance to a single bnAb, and computationally determining optimal combinations of bnAbs is an unsolved problem. Recently, machine learning models trained using resistance outcomes for multiple antibodies at once, a strategy called multi-task learning (MTL), have been shown to achieve better performance in several cases than previous approaches. We develop a new model and show that, beyond the boost in performance, MTL also helps address the previous two challenges. Specifically, we demonstrate empirically that MTL can mitigate bias from underrepresented subtypes, and that MTL allows the model to learn patterns of co-resistance between antibodies, thus providing tools to predict antibodies’ epitopes and to potentially select optimal bnAb combinations. Our analyses, publicly available at https://github.com/iaime/LBUM, can be adapted to other infectious diseases that are treated with antibody therapy.

## Introduction

Broadly neutralizing antibodies (bnAbs) exhibiting exceptional breadth and potency have revived the hope for immunotherapy against HIV-1 (1). To neutralize most viruses and to prevent viral escape, bnAbs will likely be given in combinations. For example, one cocktail, made of 3BNC117 and 10-1074, has achieved viral suppression for roughly 20 weeks without antiretroviral therapy in 9 out of 11 individuals and 13 out 17 individuals participating in two phase 1b clinical trials, respectively (2,3). Nonetheless, the general question of which bnAbs to administer together to achieve maximum efficacy is still outstanding.

Given that bnAbs target HIV’s envelope glycoprotein (Env), neutralization assays are traditionally used to determine the breadth and potency of different bnAbs against panels of Env-pseudotyped viruses (4). For each pseudovirus, these experiments determine the bnAb concentration needed to reduce infectivity by 50% or 80% (i.e., IC_50_ or IC_80_, respectively). These assays are expensive and slow. In particular, when the goal is to identify bnAbs that are likely to neutralize most viruses in a given population, there is the need for scalable computational methods to predict Env sequences’ sensitivity to bnAbs.

Several machine learning (ML) models (5–12) to map Env sequences to bnAb susceptibility have been developed using neutralization data compiled in the web server CATNAP (13). The generalizability of these methods beyond the training data is unclear, as the training datasets have HIV-1 subtype compositions that are unrepresentative of large epidemics in sub-Saharan Africa (Fig S1) (14), where the two thirds of people living with HIV-1 worldwide reside (15). This is particularly worrying since susceptibility to bnAbs can be subtype-dependent (16).

Some of the most recent ML models in predicting HIV-1 resistance to many bnAbs use multi-task learning (MTL) (9,12). The premise of MTL is that information from different but related tasks is beneficial to specific tasks of interest (17). In this context one model is trained using neutralization outcomes for multiple antibodies at once, as opposed to only considering one antibody per model. Here we show that, in addition to a boost in performance in some cases, MTL provides solutions, at least partially, to the challenges related to data imbalances and to the selection of optimal bnAb combinations.

Specifically, we empirically show that: a) MTL can mitigate bias against underrepresented HIV-1 subtypes; b) MTL allows learning patterns of co-resistance between antibodies, thus providing tools to predict antibodies’ epitopes and to potentially select optimal bnAb combinations.

## Results

### Model rationale

A common modeling choice is to align Env sequences and treat a site in the alignment as a categorical variable (5–8,10,11). However, Env is highly variable, thus making multiple sequence alignment very challenging. Natural language processing (NLP) techniques offer alignment-free methods, which leverage the distributional hypothesis originating from linguistics (18). The hypothesis stipulates that similar words tend to occur in similar contexts. This allows language models trained on large corpora to learn semantically meaningful vector representations of words, called word embeddings. In the case of modeling protein sequences, each amino acid can be treated as a word whose embedding is learned based on its co-occurrences with other amino acids in many sequences. Importantly, the embeddings need not be fixed for each amino acid, but can rather vary depending on the rest of the sequence, resulting in contextualized embeddings.

Using architectural details from (19), we trained a base Env language model to learn contextualized embeddings. This task consisted in predicting each amino acid in the sequence given the rest of the sequence. The average of these embeddings across all amino acids in a sequence can be understood as the overall vector representing the sequence. Such vectors can be used to explain variations between sequences (19). This phase of training the base model is what we call “pretraining,” which only requires Env sequences, without any neutralization data attached to them. In this work, we pretrained using 71390 Env sequences from the Los Alamos National Laboratory HIV Sequence Database (https://www.hiv.lanl.gov/). As many sequences without neutralization data are available, we hypothesized that pretraining would potentially improve the model’s generalizability, in addition to making the model learn alignment-free sequence encodings.

The second component of an input to a MTL model is the antibody of interest. Inspired by works in NLP (20–22), we represent each antibody by a unique vector. We call this vector an antibody context. Based on the distributional hypothesis, we reasoned that differences between learned antibody contexts would encode correlations between antibodies’ resistance profiles, thus offering insights into potential optimal bnAb combinations. For simplicity, we did not consider antibody sequences themselves, unlike in (12). Instead, antibody contexts were randomly initialized and tuned using neutralization data linking antibodies to Env sequences in the training data. The resulting MTL model is what we refer to as a language-based universal model (LBUM) (Fig 1). Further details of the model are given in *Methods*.

**Fig 1.**
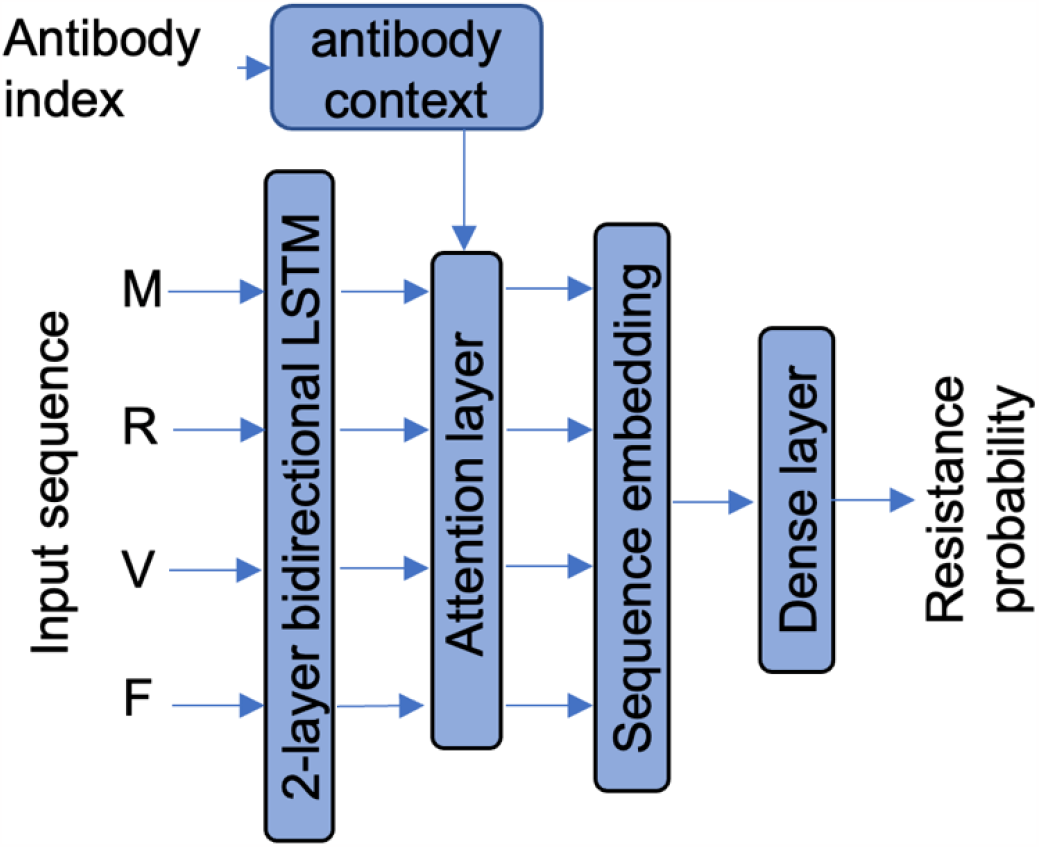
The architecture of a language-based universal model (LBUM).

### No single model dominates across all bnAbs

We considered 33 bnAbs targeting five different epitopes: the membrane-proximal external region (MPER), the gp120-gp41 interface, the CD4 binding sites (CD4bs), the third constant region and the third variable loop (C3/V3), and the first and second variable loops (V1/V2) (16,23). The first two columns of Table 1 show the epitope that each of the 33 bnAbs targets.

**Table 1.** Models’ AUC. Area under the receiver operating characteristic curve (AUC). GBM is Gradient Boosting Machines; RF is Random Forests; LBUM is language-based universal model; ENS is the ensemble model that averages predictions from GBM, RF and LBUM. The red shade means that LBUM had a better score than both RF and GBM models did. The blue shade means the ensemble model scored better than all three individual models did. Numbers between parentheses are standard deviations.

| Epitope | BnAb | GBM | RF | LBUM | ENS |
| --- | --- | --- | --- | --- | --- |
| CD4bs | VRC07 | 0.78 (0.08) | 0.85 (0.03) | 0.93 (0.03) | 0.94 (0.05) |
|  | VRC01 | 0.86 (0.03) | 0.89 (0.02) | 0.91 (0.01) | 0.93 (0.03) |
|  | NIH45-46 | 0.84 (0.04) | 0.84 (0.05) | 0.91 (0.03) | 0.93 (0.03) |
|  | VRC-CH31 | 0.77 (0.06) | 0.78 (0.07) | 0.82 (0.10) | 0.86 (0.06) |
|  | VRC-PG04 | 0.80 (0.10) | 0.82 (0.07) | 0.85 (0.07) | 0.89 (0.08) |
|  | HJ16 | 0.53 (0.07) | 0.53 (0.07) | 0.63 (0.04) | 0.62 (0.07) |
|  | 3BNC117 | 0.89 (0.04) | 0.92 (0.04) | 0.91 (0.02) | 0.93 (0.03) |
|  | VRC03 | 0.83 (0.03) | 0.83 (0.01) | 0.81 (0.05) | 0.86 (0.02) |
|  | VRC13 | 0.83 (0.01) | 0.86 (0.03) | 0.75 (0.07) | 0.87 (0.02) |
|  | b12 | 0.80 (0.04) | 0.81 (0.03) | 0.73 (0.05) | 0.81 (0.03) |
| C3/V3 | DH270.1 | 0.89 (0.03) | 0.91 (0.05) | 0.94 (0.02) | 0.95 (0.01) |
|  | VRC29.03 | 0.78 (0.10) | 0.80 (0.07) | 0.86 (0.06) | 0.87 (0.07) |
|  | DH270.5 | 0.92 (0.02) | 0.93 (0.02) | 0.93 (0.03) | 0.96 (0.02) |
|  | DH270.6 | 0.93 (0.05) | 0.95 (0.03) | 0.93 (0.03) | 0.97 (0.02) |
|  | PGT135 | 0.77 (0.05) | 0.77 (0.06) | 0.76 (0.04) | 0.79 (0.04) |
|  | PGT128 | 0.87 (0.04) | 0.86 (0.02) | 0.84 (0.03) | 0.89 (0.02) |
|  | 2G12 | 0.91 (0.02) | 0.90 (0.03) | 0.85 (0.03) | 0.90 (0.03) |
|  | PGT121 | 0.91 (0.02) | 0.91 (0.02) | 0.84 (0.02) | 0.93 (0.02) |
|  | 10-1074 | 0.97 (0.02) | 0.97 (0.01) | 0.88 (0.04) | 0.98 (0.01) |
| MPER | 4E10 | 0.60 (0.12) | 0.63 (0.15) | 0.73 (0.06) | 0.72 (0.10) |
|  | 2F5 | 0.96 (0.03) | 0.95 (0.03) | 0.90 (0.03) | 0.95 (0.03) |
| gp120-gp41 | 35O22 | 0.56 (0.13) | 0.62 (0.08) | 0.59 (0.06) | 0.61 (0.10) |
|  | VRC34.01 | 0.82 (0.03) | 0.80 (0.07) | 0.74 (0.06) | 0.81 (0.04) |
|  | PGT151 | 0.78 (0.06) | 0.79 (0.04) | 0.65 (0.07) | 0.78 (0.05) |
|  | 8ANC195 | 0.88 (0.05) | 0.86 (0.05) | 0.63 (0.04) | 0.88 (0.05) |
| V1/V2 | VRC26.08 | 0.88 (0.02) | 0.87 (0.02) | 0.93 (0.03) | 0.94 (0.02) |
|  | VRC26.25 | 0.86 (0.03) | 0.85 (0.03) | 0.87 (0.06) | 0.91 (0.02) |
|  | PG16 | 0.82 (0.04) | 0.83 (0.03) | 0.84 (0.02) | 0.85 (0.03) |
|  | PG9 | 0.85 (0.03) | 0.84 (0.02) | 0.85 (0.03) | 0.89 (0.03) |
|  | CH01 | 0.79 (0.03) | 0.79 (0.03) | 0.76 (0.05) | 0.82 (0.02) |
|  | PGDM1400 | 0.90 (0.02) | 0.89 (0.02) | 0.90 (0.02) | 0.94 (0.02) |
|  | VRC38.01 | 0.69 (0.10) | 0.66 (0.03) | 0.66 (0.09) | 0.71 (0.09) |
|  | PGT145 | 0.85 (0.03) | 0.85 (0.05) | 0.79 (0.07) | 0.87 (0.03) |

Our aim was not to simply develop models that predict HIV-1 resistance to the 33 bnAbs; however, we still compared the LBUM to models developed with classical machine learning algorithms, namely random forests (RF) and gradient boosting machines (GBM). We caution against comparisons to previously published performances since CATNAP data has changed over time, and preprocessing and model-selection techniques vary across publications (24).

We assessed models using three metrics: the area under the receiver operating characteristic curve (AUC), interpreted as the probability that a model ranks resistant sequences above sensitive ones; the area under the precision-recall curve (PR AUC), which measures how the model trades off precision for sensitivity, an important metric especially when resistance sequences are rare; and the binary cross-entropy (Log Loss), which measures the difference between predicted resistance probabilities and the ground truth.

The LBUM achieved higher AUC and higher PR AUC than both RF and GBM models did on the same 12 bnAbs out of the 33 bnAbs (Table 1, S1 Table). Notably, 6 of those 12 bnAbs target the CD4bs. In terms of Log Loss, the LBUM scored better than both RF and GBM models did on 10 bnAbs, 4 of which target the CD4bs (S1 Table). Interestingly, the LBUM’s AUC and PR AUC on two of the 10 bnAbs (i.e., PGT135 and PGT128) was slightly lower than RF and GBM models’. Otherwise, for the remaining 8 bnAbs, the lower LBUM’s Log Loss meant higher AUC and higher PR AUC than RF and GBM models’.

Overall, there was no single model that consistently outperformed all other models across all bnAbs. Nevertheless, averaging predicted resistance probabilities from the three models—defining the ensemble model, ENS—mitigated underperformances from individual models. The ensemble model achieved the highest AUC on 24 bnAbs (Table 1), the highest PR AUC on 21 bnAbs (S1 Table), and the lowest Log Loss on 22 bnAbs (S1 Table).

### Multi-task learning can mitigate HIV-1 subtype bias

Publicly available training datasets are very imbalanced in terms of HIV-1 subtypes (Fig S1), which can compromise models’ generalizability to underrepresented subtypes, a problem we call “subtype bias” hereafter. The LBUM uses, in addition to the usual bnAb data, large numbers of Env sequences with no neutralization data, and also data from antibodies not deemed as bnAbs. We hypothesized that both of these data sources help to mitigate subtype bias, because they have more balanced availability across subtypes than bnAb data. To test the two aspects separately, one would ideally vary the composition of subtypes at different training stages. However, only subtype B and subtype C had sufficient data to run meaningful tests (Fig S1). In addition, data on bnAbs NIH45-46, VRC13, VRC07, and VRC26.08 was excluded from this analysis because it lacked enough subtype and phenotype diversity.

To quantify the level of subtype bias, we trained two models, one with only subtype B data and one with only subtype C data. We then evaluated the models on the subtype B and subtype C bnAb testing sets separately. With the exception of gp120-gp41 bnAbs (n=4) and MPER bnAbs (n=2), the AUC was greater by roughly 0.3 on the matched subtype than on the unmatched (Fig 2A). The model trained on both subtypes did equally well at classifying both subtypes (Fig 2A). While experimental data from antibody neutralization assays was not available for all subtypes, there was sequence data for all subtypes. To test whether using the additional sequence data improved the generalizability of the models, we re-trained the subtype-specific model but included both subtypes in the initial pretraining step. Unfortunately, using the additional sequences improved the generalizability only minimally, if at all (Fig 2B).

**Fig 2.**
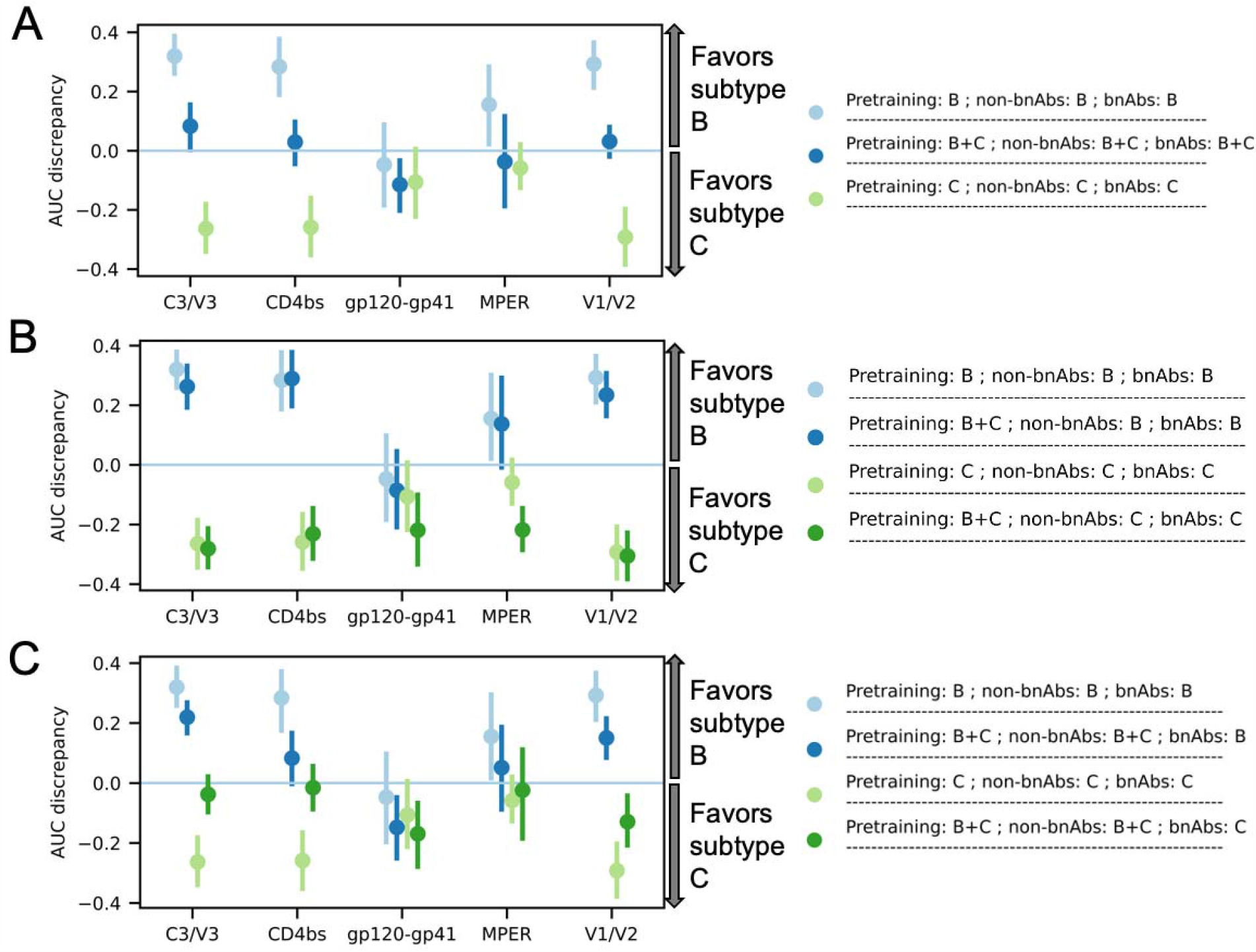
Effect of subtype representativeness on AUC. We named models according to the subtype combinations contained in the pretraining data (shown as “Pretraining”), in data on non-bnAbs (shown as “non-bnAbs”), and in bnAb data (shown as “bnAbs”). AUC discrepancy means AUC on subtype B minus AUC on subtype C. (A) shows the bias introduced by only training on one subtype, and how that bias is eliminated by more subtype diversity. (B) shows that subtype representativeness in the pretraining data reduces subtype bias only to a small extent, if at all. (C) shows how subtype representativeness in non-bnAb data reduces subtype bias. Error bars represent the 95% confidence intervals computed using 1000 bootstrap samples.

While neutralization data for bnAbs is limited, there is often data for other antibodies, which we label as “non-bnAbs”. We tested whether including this non-bnAb data in the training of the subtype-specific models improved their generalizability. Except on gp120-gp41 bnAbs, these models showed much greater generalizability, with the difference in AUC between the two subtypes dropping to roughly 0.1 or less in many cases (Fig 2C).

PR AUC and Log Loss generally showed similar patterns of subtype bias to those seen for AUC (Fig S2 and Fig S3). That is, subtype representativeness in non-bnAb data improved PR AUC on the unmatched subtype (Fig S2C & Fig S3C), while pretraining with the subtype of interest had very minimal effects on subtype bias (Fig S2B & Fig S3B). The exceptions for gp120-gp41 and MPER bnAbs remained. We also note that Log Loss discrepancy did not change as much on C3/V3 bnAbs for some models, despite subtype representativeness in non-bnAb data (Fig S3C).

### Do the learned antibody contexts encode co-resistance patterns?

If learned antibody contexts encode co-resistance patterns, we would expect many bnAbs targeting similar epitopes to have similar contexts, given that bnAbs targeting similar epitopes tend to have similar resistance patterns (16). Clustering by epitope could be observed after projecting the dimensionality of the antibody contexts to a two-dimensional space (Fig 3A-E). Without any further training we could predict bnAb epitopes solely based on the epitope targeted by the closest bnAb in that context space with at least 72% accuracy (Table 2). We defined closeness between bnAbs in terms of cosine similarity, L1 distance and L2 distance between their context vectors. In at least 85% of cases, at least one of the 5 closest bnAbs targeted the same epitope as the bnAb in question (Table 2), further suggesting that antibody contexts captured epitope-specific resistance patterns.

**Table 2.** Proportion of bnAbs whose epitope was targeted by at least one of the closest bnAbs. The numbers between parentheses are standard deviations, since there are 5 LBUMs that resulted from performing 5-fold cross-validation.

| Number of closest bnAbs considered | Cosine similarity | L1 distance | L2 distance |
| --- | --- | --- | --- |
| 1 | 0.74 (0.06) | 0.72 (0.04) | 0.75 (0.05) |
| 2 | 0.82 (0.03) | 0.79 (0.06) | 0.79 (0.06) |
| 3 | 0.89 (0.04) | 0.83 (0.06) | 0.84 (0.03) |
| 4 | 0.94 (0.02) | 0.84 (0.06) | 0.84 (0.04) |
| 5 | 0.95 (0.02) | 0.85 (0.05) | 0.87 (0.05) |

**Fig 3.**
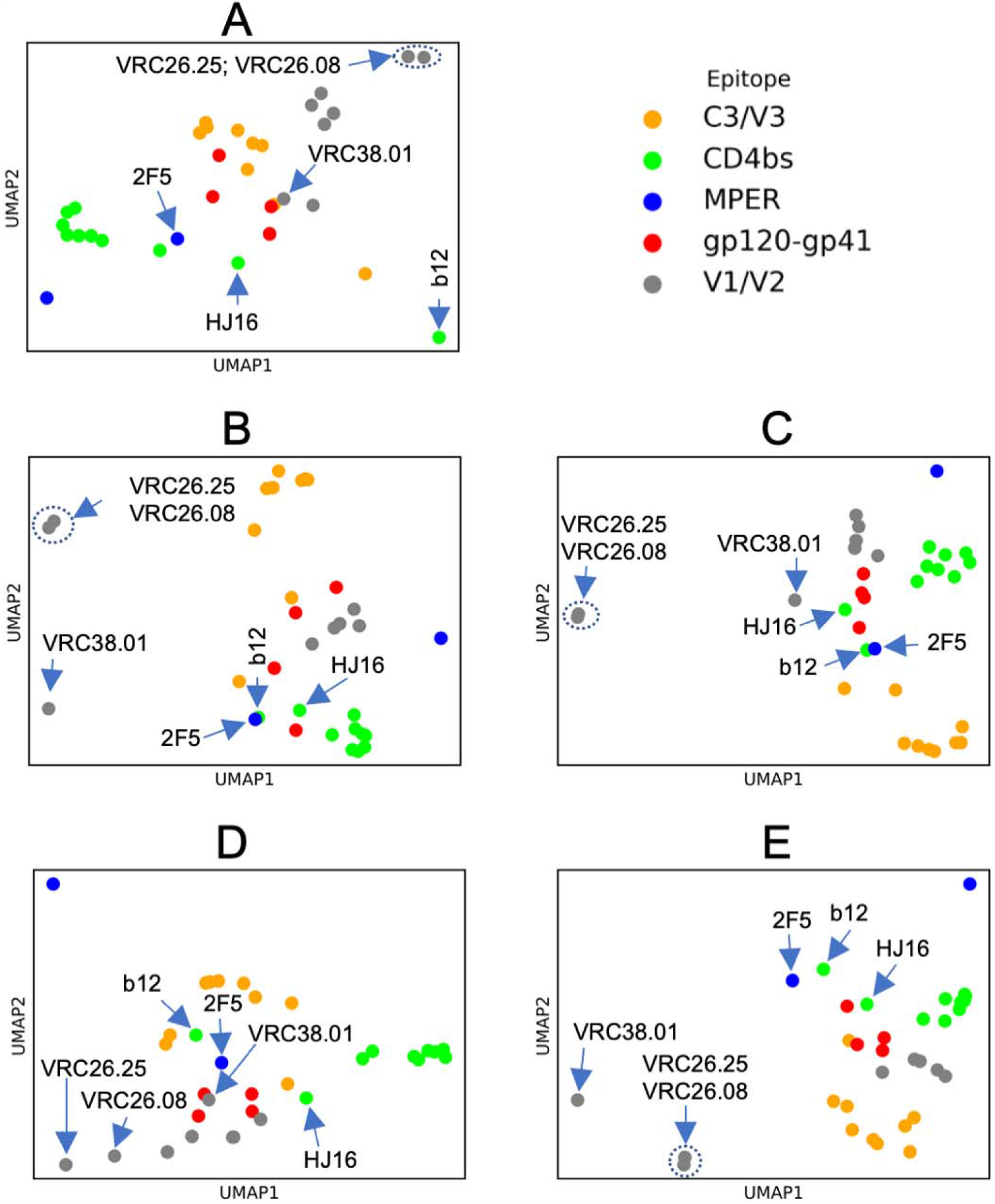
Learned antibody contexts. Antibody contexts (i.e., vector representations of antibodies) learned as part of the attention mechanism. As a result of performing 5-fold cross-validation, 5 LBUMs were available, and antibody contexts from each of these are shown in one of the 5 subfigures. BnAbs are color-coded according to their epitopes. Arrows point to some bnAbs of interest; dashed-line circles show where one arrow points to two very close bnAbs.

Although bnAbs targeting similar epitopes generally tend to have similar resistance profiles, that is not always the case. Indeed, we observed outliers in epitope clusters (Fig 3). We therefore investigated whether such within-epitope dissimilarities imply different resistance patterns among bnAbs targeting similar epitopes. A known example of dissimilar patterns within the V1/V2 epitope was captured by learned bnAb contexts: VRC26.08 and VRC26.25 clustered away from the rest of V1/V2 bnAbs (Fig 3A,B,C,E). Contrary to the rest of V1/V2 bnAbs, the potency of CAP256-VRC26 bnAbs, which include the two bnAbs, is known to be inversely dependent on the presence of a glycan at the N160 position in Env (25). VRC38.01 was also another outlier in the V1/V2 cluster (Fig 3B,C&E). Unlike the other V1/V2 bnAbs, VRC38.01 has a unique binding mode that allows it to have non-protruding heavy-chain complementarity-determining region 3 (HCDR3) (26). Furthermore, b12 and HJ16 clustered away from the rest of CD4bs bnAbs (Fig 3A-D). Unlike VRC01-like bnAbs, b12 does not mimic CD4 binding and binds Env in its relaxed conformation (27). On the other hand, HJ16 interferes with CD4 by binding to a glycan at site N270 in Env. The mutation N270D, which removes the glycan, makes viruses resistant to HJ16, while making them more sensitive to other CD4bs bnAbs such as VRC01 and VRC03 (28). Another interesting dissimilarity was between 2F5 and 4E10, which target different MPER regions (29). Mutations with opposing effects on resistances to 2F5 versus 4E10 have been reported (16).

Finally, we note that some bnAbs, such as 2F5 and b12 (Fig 3B&C), appeared to have similar bnAb contexts despite targeting different epitopes. Whether such cases imply cross-epitope resistance correlation is an interesting question, which we leave for future work.

## Discussion

In summary, we developed a model to predict the neutralization of different HIV-1 Env sequences by different broadly neutralizing antibodies (bnAbs). Our model, which we named a language-based universal model (LBUM), is a type of multi-task learning (MTL) model. The LBUM was pretrained using Env sequences with no associated neutralization data and fine-tuned with Env sequences with both non-bnAb and bnAb outcome data. We first showed that the LBUM’s performance is comparable to that of Gradient Boosting Machine (GBM) models and Random Forest (RF) models, with some improvements over both methods (Table 1, S1 Table). Unlike the other two methods, the LBUM does not require aligning input Env sequences, which is an advantage given the incredible variability of Env, including structural variability that makes alignment challenging. As for previous methods, all models in this work were trained to predict *in vitro* bnAb resistance: we did not validate them with clinical outcomes, and we relied on data showing correlations between *in vitro* susceptibility to bnAbs and *in vivo* outcomes (30).

A thorough systematic comparison between published methods requires testing different combinations of preprocessing techniques, feature selection methods and learning algorithms. In this work, we only compared learning algorithms applied on full Env sequences with neutralization data preprocessed similarly. We compared the LBUM to both RF and GBM models because both boosting trees and RF underlie recently published methods that do not use neural networks (5,7).

The most common subtypes in sub-Saharan Africa are A, C, D and several circulating recombinant forms (CRFs) (14). CATNAP, from which most of the training datasets come, has mostly subtype B and subtype C sequences (Fig S1). This subtype mismatch is problematic because, as we have shown, models do not necessarily generalize across subtypes (Fig 2A, Fig S2A, Fig S3A). MTL and pretraining give access to more data with potentially more subtype representativeness. Although no solution trumps having all subtypes represented in bnAb data, our results suggest that MTL can alleviate subtype bias if neutralization data with all subtypes is available for antibodies not considered bnAbs (Fig 2B, Fig S2B, Fig S3B).

We introduced the concept of antibody contexts, which we defined as vector representations unique to each antibody and updated during the fine-tuning process. We showed that bnAbs targeting similar epitopes tended to have similar contexts, to such an extent that we could use closeness between antibody contexts to predict antibody epitopes (Table 2). In this regard, our methods can be used to generate hypotheses about epitopes targeted by new antibodies, as long as relevant neutralization data is part of the LBUM’s training data. Nonetheless, some bnAbs had distant contexts despite targeting similar epitopes (Fig 3), and we highlighted known mechanistic reasons that support our hypothesis that differences in antibody contexts capture differences in resistance profiles. A possible limitation is that negatively correlated resistance profiles can possibly lead to similar bnAb contexts, the same way antonyms can have similar word embeddings in the English language (31). Nonetheless, we assumed that such cases were rare if present at all, given that most bnAb contexts tended to cluster per epitope. Analyses of structural data on antibodies and their respective targets on Env will help further show the extent to which antibody contexts capture co-resistance patterns.

An interesting extension of our methods could be to pretrain using generic protein language models, such as those in the BERT and ESM families (32,33). We expect MTL models’ performance to increase as their size increases along with the increase in the quantity and diversity of their training data. We chose small architectures because of computational requirements imposed by deep neural network models and because of the availability of only small amounts of data on which to fine-tune.

The potential of MTL revealed in our study addresses key challenges in HIV-1 vaccine research. All models developed in this work, along with the used code, can be found at https://github.com/iaime/LBUM. The framework presented here is a starting point towards designing effective immunotherapies. We hope that our analyses can be relevant to other infectious diseases for which monoclonal antibodies are being explored as therapeutic solutions.

## Materials and methods

### Data preprocessing

We binarized the neutralization outcome—resistant or sensitive—i.e., we aimed to predict whether positive neutralization is observed within a certain range of antibody concentrations. This is because the main use-case envisioned for our models is the identification of bnAbs that are likely to neutralize most viruses in given populations. Once bnAbs with largest coverage are identified, other methods will need to be used to determine the bnAbs’ exact potencies.

We determined the phenotype based on IC_50_ because this was more commonly available than IC_80_ for Env sequences in CATNAP. We transformed left-censored IC_50_ values to the detection threshold. That is, <x values became x. Since CATNAP Env sequences could have multiple IC_50_ values from different studies, we calculated the geometric mean whenever more than one value was available, as long as none of the values was right-censored. If any reported IC_50_ value for a sequence-antibody pair was right-censored, the sequence was deemed resistant to the antibody, unless the detection threshold was lower than 10 μg/mL, in which case the sequence-antibody entry was discarded. If no right-censored IC_50_ values were recorded for the sequence-antibody entry and if the geometric mean IC_50_ was greater than 50 μg/mL, the sequence was also labeled as resistant to the antibody. Otherwise, the sequence was labeled as sensitive to that antibody (i.e., sensitive sequences had no right-censored IC_50_ values and the geometric mean IC_50_ was less than 50 μg/mL).

In all our experiments, we only considered sequences that are 800 to 900 amino acid long (ignoring non-amino-acid characters) to match the expected length of a full Env sequence. Part of our analysis compared our model to random forests (RF) and gradient boosting machines (GBM) models. Since RF and GBM models required aligned sequences, we used the alignment provided in CATNAP to one-hot encode sequences. That is, each amino acid was represented as a vector of all zeros except a 1 at the index of that amino acid. For non-amino acid characters, the vector was all zeros. The LBUM did not require aligning sequences, and all non-amino acid characters were removed from their input sequences.

### Gradient Boosting Machines and Random Forests

Both GBM and RF build ensemble models based on decision trees. For complete mathematical descriptions of GBM and RF, we refer to (34) and (35), respectively. 5-fold nested cross-validation was used to select and evaluate both types of models. Log Loss was used to select the best classifiers. Table 3 provides all hyperparameters considered. Both GBM and RF models were implemented using scikit-learn (v1.1.1) (36).

**Table 3.** Hyperparameters considered in the development of different models.

| Model type | Hyperparameters |
| --- | --- |
| Random Forests | max depth: 1, 2, 3, 4, 5<br>max features: 0.03, 0.1, 0.2, 0.3, 0.5<br>number of trees: 10, 50, 100, 500, 1000<br>calibration method: “sigmoid”, “isotonic” |
| Gradient Boosting Machines | learning rate: 0.001, 0.01, 0.05, 0.1, 0.2<br>max features: 0.03, 0.1, 0.2, 0.3, 0.5<br>max depth: 1, 2, 3, 4, 5<br>number of trees: 10, 50, 100, 500, 1000<br>calibration method: “sigmoid”, “isotonic” |
| Language-based universal model | learning rate: 0.0001, 0.0003, 0.001, 0.003<br>antibody context dimension: 32, 64, 128, 256<br>dropout rate: 0.1, 0.2, 0.3, 0.4, 0.5<br>number of pretrained layers to unfreeze: 0, 2, 4, all<br>classification loss weight: 0.5, 0.6, 0.7, 0.8, 0.9, 1 |

### Language-based universal model (LBUM)

The overall architecture of the proposed LBUM is illustrated in Fig 1 and rationalized in the *Results* section. There were two main steps in the development of the LBUM, namely pretraining and fine-tuning. First, we pretrained the model using the same architectural and training details as Hie *et al*’s method (19). Essentially, the method was a two-layer bidirectional Long Short-Term Memory (LSTM) model (37). Next, we finetuned the LBUM using data on 378 antibodies in addition to the 33 bnAbs of interest. During fine-tuning, we froze all pretrained layers except the last right and left LSTM layers, and we applied early stopping with a 10-epoch patience.

As a regularization technique, we added a secondary output layer in the LBUM that directly predicts log_10_(IC_50_). However, sequence-antibody pairs with IC_50_ beyond the detection threshold (i.e., right-censored IC_50_) did not contribute towards the training of the regression branch. A question not addressed here is how to incorporate censored data into the training data of models that predict IC_50_. For now, we recommend against making predictions with the regression branch of the trained model, as it cannot be relied on given its biased training data. The LBUM’s overall loss function was simply the weighted average of binary cross-entropy and mean squared error, with weights of 0.6 and 0.4, respectively.

The dimension of antibody context vectors was set to 128. These vectors were finetuned through an attention mechanism that was a combination of at least three methods (20–22). To visualize the antibody contexts in Fig 3, we used the Uniform Manifold Approximation and Projection algorithm (UMAP) (38). Below we detail the attention layer.

Let *C*_*t*_ be the context of an antibody t. Let *E*_*j*_ be the embedding of a token *j* in a sequence x of length *n*. The attention weight *a*_*j*_ to the token <u>j</u> given the antibody *t* context was calculated as follows:

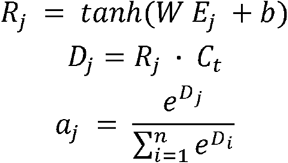

where *W* and are weight matrix and bias vector, respectively, and *tann* is the hyperbolic tangent used as an activation function. We note that 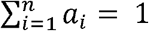 for each sequence. The weighted average embedding 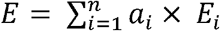 was then input to a dense output layer. We added two dropout layers at a 10% rate, one before the attention layer and another before the final dense layer. We also used temperature scaling as the calibration method (39). Each antibody had its own temperature value initialized to 1.5 and optimized during the fine-tuning process. At inference time, we averaged predictions from running 10 forward passes with dropout turned on.

The LBUM hyperparameters were tuned on non-bnAb data to avoid data leakage. Using Bayesian optimization implemented in KeraTuner (40), we tuned the learning rate for the fine-tuning phase, the dimension of antibody context vectors, the dropout rate, the number of pretrained layers to unfreeze during fine-tuning, and the weights for the classification and regression output branches of the LBUM. Considered values for these hyperparameters are shown in Table 3. Values that achieved the lowest binary crossentropy within 10 trials were chosen for the final model. All the other hyperparameters of the LBUM were set to default values in Tensorflow Keras (v2.12.0).

## Acknowledgments

We thank all members of the Pathogen Dynamics Group in the Big Data institute at the University of Oxford for their helpful feedback during the course of this project. This work was supported by a Li Ka Shing Foundation Grant awarded to C.F. The computational aspects of this research were funded from the NIHR Oxford BRC with additional support from the Wellcome Trust Core Award Grant Number 203141/Z/16/Z. The views expressed are those of the author(s) and not necessarily those of the NHS, the NIHR or the Department of Health.

## Notes

https://github.com/iaime/LBUM

### Competing Interest Statement

The authors have declared no competing interest.

