## Supporting information for "Learning patterns of HIV-1 co-resistance to broadly neutralizing antibodies with reduced subtype bias using multi-task learning"

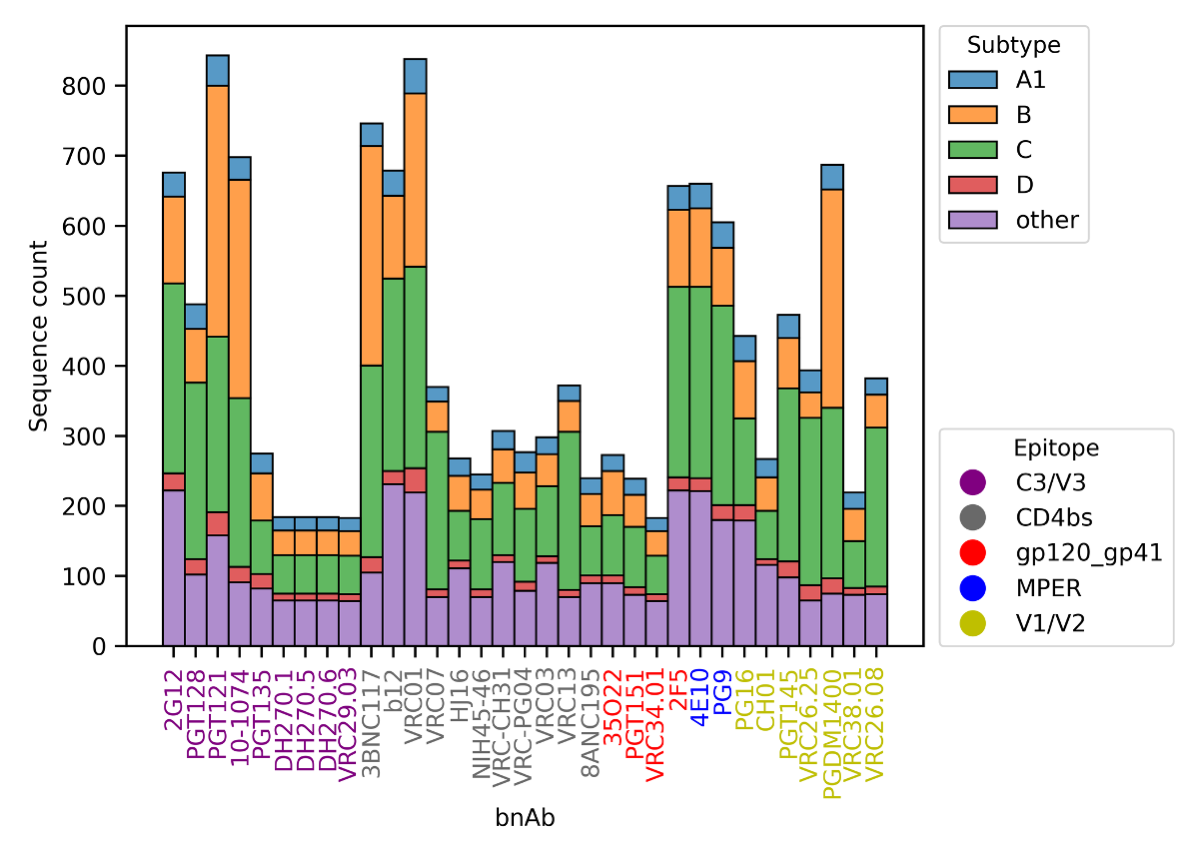

**Fig S1. CATNAP data.** Shown are the counts of Env sequences for which neutralization assay data is available for each of the 33 bnAbs we considered. Distributions per subtype are color-coded. BnAbs are also color-coded according to the epitope they target.

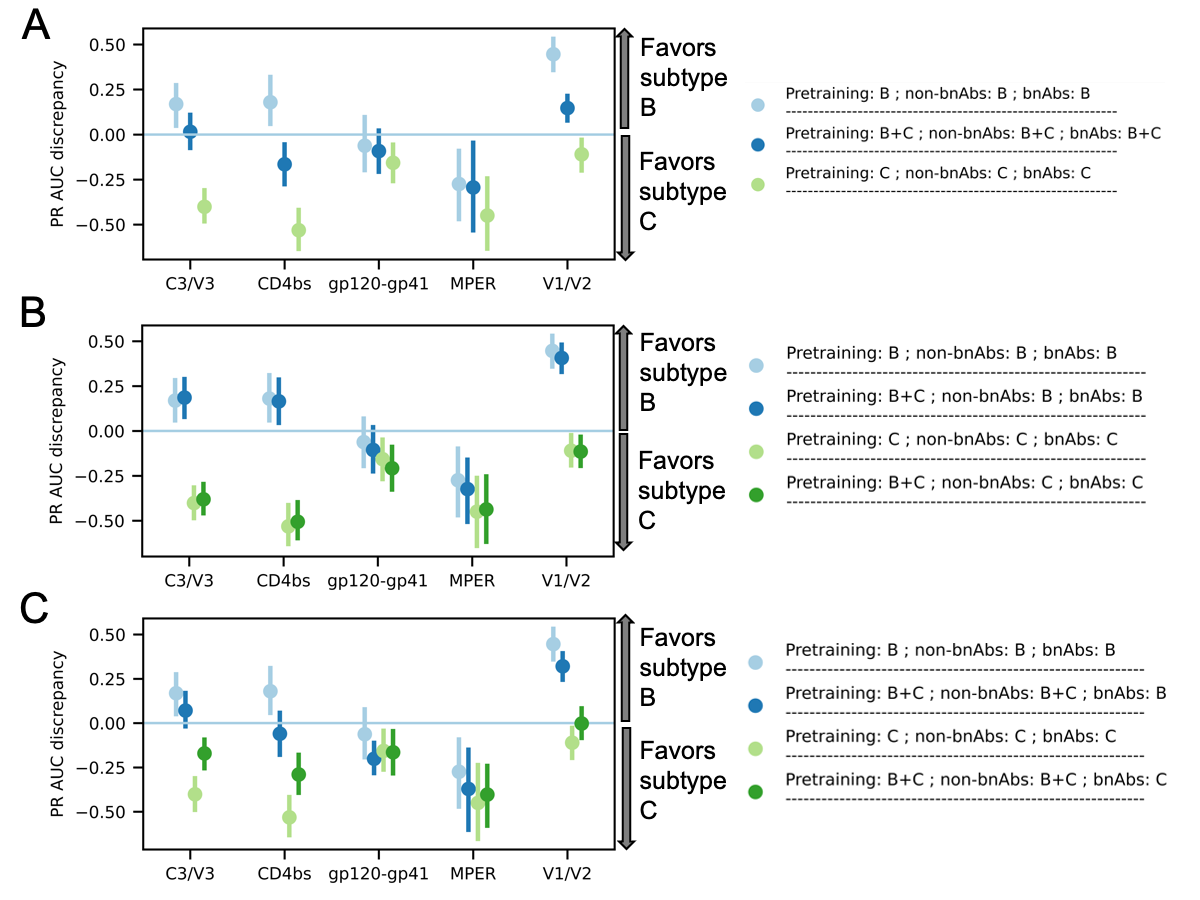

**Fig S2. Effect of subtype representativeness on PR AUC.** We named models according to subtype combinations contained in the pretraining data (shown as “Pretraining”), in data on non-bnAbs (shown as “non-bnAbs”), and in bnAb data (shown as “bnAbs”). PR AUC discrepancy means PR AUC on subtype B minus PR AUC on subtype C. (A) shows the bias introduced by only training on one subtype, and how that bias is eliminated by more subtype diversity. (B) shows that subtype representativeness in the pretraining data reduces subtype bias only to a small extent, if at all. (C) shows how subtype representativeness in non-bnAb data reduces subtype bias. Error bars represent the 95% confidence intervals computed using 1000 bootstrap samples.

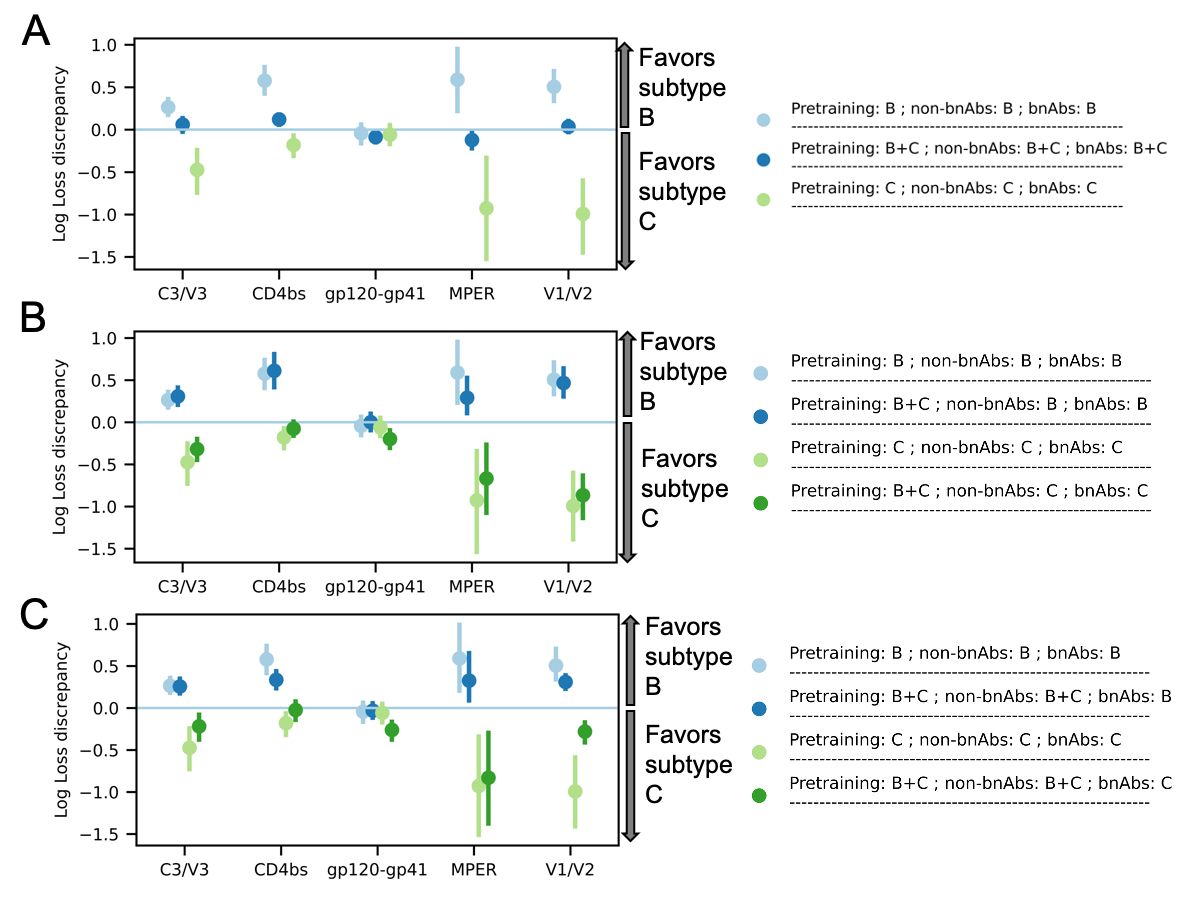

**Fig S3. Effect of subtype representativeness on Log Loss.** We named models according to subtype combinations contained in the pretraining data (shown as “Pretraining”), in data on non-bnAbs (shown as “non-bnAbs”), and in bnAb data (shown as “bnAbs”). Log Loss discrepancy means Log Loss on subtype C minus Log Loss on subtype B. (A) shows the bias introduced by only training on one subtype, and how that bias is eliminated by more subtype diversity. (B) shows that subtype representativeness in the pretraining data reduces subtype bias only to a small extent, if at all. (C) shows how subtype representativeness in non-bnAb data reduces subtype bias. Error bars represent the 95% confidence intervals computed using 1000 bootstrap samples.

**S1 Table. Models’ PR AUC and Log Loss.** The area under the precision-recall curve (PR AUC) and the binary cross-entropy (Log Loss) are reported. GBM is Gradient Boosting Machines; RF is Random Forests; LBUM is language-based universal model; ENS is the ensemble model that averages predictions from GBM, RF and LBUM. The red shade means that LBUM had a better score than both RF and GBM models did. The blue shade means the ensemble model scored better than all three individual models did. Numbers between parentheses are standard deviations.

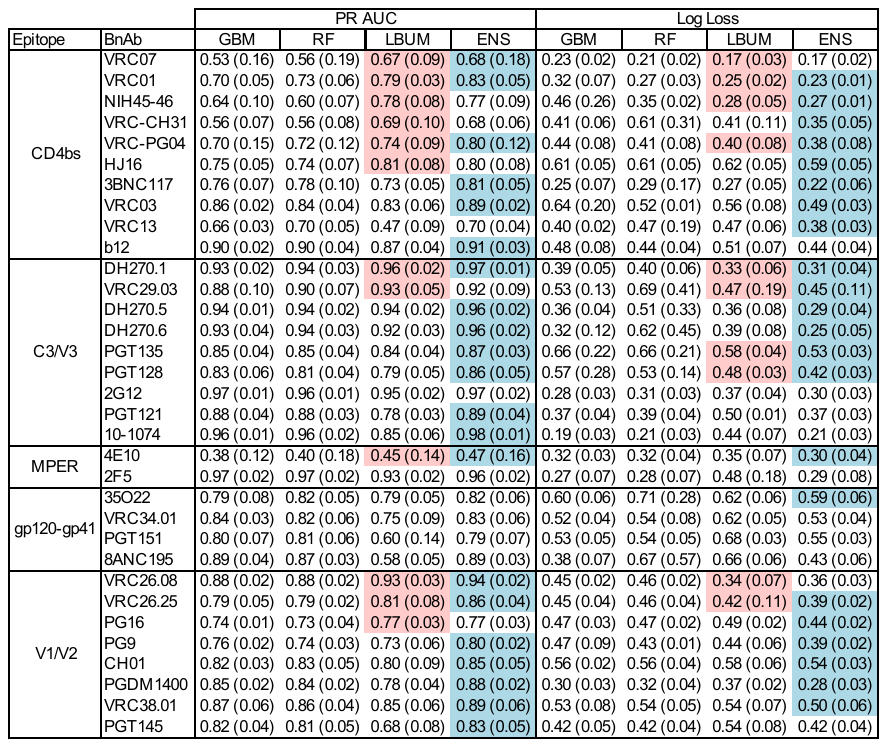
